# Copy number variation and neurodevelopmental problems in females and males in the general population

**DOI:** 10.1101/236042

**Authors:** Joanna Martin, Kristiina Tammimies, Robert Karlsson, Yi Lu, Henrik Larsson, Paul Lichtenstein, Patrik Magnusson

**Affiliations:** Drs Martin, Karlsson, Lu, Larsson, Lichtenstein and Magnusson are at the Department of Medical Epidemiology & Biostatistics, Karolinska Institutet, Stockholm, Sweden.; Dr Martin is also at the MRC Centre for Neuropsychiatric Genetics and Genomics, Cardiff University, Cardiff, UK.; Dr Tammimies is at the Center of Neurodevelopmental Disorders at Karolinska Institutet (KIND), Department of Women’s and Children’s Health, Karolinska Institutet and Center for Psychiatry Research, Stockholm, Sweden.; Dr Larsson is also at the School of Medical Sciences, Örebro University, Örebro, Sweden.

**Keywords:** Neurodevelopmental disorders, Copy number variation, Genetics, ADHD, Autism

## Abstract

**Objective:** Neurodevelopmental problems (NPs) are childhood phenotypes that are more common in males. Conversely, anxiety and depression (which are frequently comorbid with NPs) are more common in females. Rare copy number variants (CNVs) have been implicated in clinically-defined NPs. Here, we aimed to characterise the relationship between rare CNVs with NPs and anxiety/depression in a population sample of twin children. Additionally, we examined whether sex-specific CNV effects underlie the sex bias of these disorders.

**Method:** We analysed a sample of N=12,982 children, of whom 5.3% had narrowly-defined NPs (clinically-diagnosed), 20.9% had broadly-defined NPs (based on validated screening measures, but no diagnosis) and 3.0% had clinically-diagnosed anxiety or depression. Rare (<1% frequency) CNVs were categorised by size (medium: 100-500kb or large: >500kb), type (duplication or deletion) and putative relevance to NPs (affecting previously implicated loci or evolutionarily-constrained genes). We tested for associations between the different CNV categories with NPs and anxiety/depression, followed by examination of sex-specific effects.

**Results:** Medium deletions (OR(CI)=1.18(1.05-1.33),p=0.0053) and large duplications (OR(CI)=1.45(1.19-1.75),p=0.00017) were associated with broadly-defined NPs. Large deletions (OR(CI)=1.85(1.14-3.01),p=0.013) were associated with narrowly-defined NPs. The effect sizes increased for large NP-relevant CNVs (broadly-defined: OR(CI)=1.60(1.06-2.42),p=0.025; narrowly-defined: OR(CI)=3.64(2.16-6.13),p=1.2E-6). No sex differences in CNV burden were found in individuals with NPs (p>0.05). In individuals diagnosed with anxiety or depression, females were more likely to have large CNVs (OR(CI)=3.75(1.45-9.68),p=0.0064).

**Conclusion:** Rare CNVs are significantly associated with both narrowly- and broadly-defined NPs in a general population sample of children. Our results also suggest that large, rare CNVs may show sex-specific phenotypic effects.

## Introduction

Neurodevelopmental problems (NPs) are psychiatric and cognitive phenotypes that are characterised by childhood onset, high levels of comorbidity with other NPs and other psychiatric conditions, and being more commonly diagnosed in males than females^1^. Common NPs include attention-deficit/hyperactivity disorder (ADHD), autism spectrum disorder (ASD), motor problems, tic disorders, and learning difficulties. NPs are highly heritable, with a complex genetic architecture according to emerging genetic studies^2-4^. Rare genetic variants, such as copy number variants (CNVs) and protein-truncating variants have been implicated in ASD, intellectual disability (ID), ADHD, developmental coordination disorder and Tourette’s syndrome^5-11^. In contrast, the contribution of rare variants to risk of childhood anxiety and depression has yet to be investigated.

Although genetic studies have tended to focus on categorically-defined clinical neurodevelopmental disorders, evidence is emerging to suggest that these disorders reflect a quantitative extreme of continuously-distributed traits in the general population. Twin studies suggest that ADHD, ASD and mild ID share genetic risks with related traits in the population^12^. Genome-wide studies further demonstrate that common variants are shared to a degree between disorders and traits related to ADHD, ASD and Tourette’s syndrome^13-15^.

Evidence for shared rare variants across disorders and traits is more limited. In a study of autism simplex families, adaptive and social-communication difficulties in ASD probands and their unaffected siblings were associated with very rare *de novo* protein-truncating variants, with no qualitative distinction between probands and siblings in this association^14^. A few studies have examined the manifestation of CNVs in the general population, focusing on ‘neuropsychiatric’ CNVs (i.e. loci robustly implicated in psychiatric disorders). These neuropsychiatric CNVs were associated with lower general assessment of function, more depression, a history of reading and mathematical learning difficulties, lower cognitive abilities, and educational attainment in adult population controls^16-18^. It appears that the same rare CNV loci are not only associated with multiple clinically-diagnosed disorders (e.g. ASD, ID, schizophrenia and ADHD^5,19,20^) but also psychiatric and cognitive outcomes in adults in the general population^16-18^. The manifestation of CNVs in relation to more broadly-defined NPs in a childhood general population sample has yet to be examined.

One prominent characteristic of NPs is that they are more commonly diagnosed in males than females^1^. Population studies of childhood NPs also suggest that males score higher on measures of NPs than females^21-23^. Although the reasons for this are still unclear, a few possible explanations have been proposed. First, it has been suggested that females are protected (e.g. by hormonal differences) from manifesting NPs and require a higher burden of risk to develop NPs. Evidence from several family studies seems to support this hypothesis^23-29^, albeit other studies are inconsiste^30-33^. Evidence from common variant analyses is also mixed, with the largest studies not finding differences between males and females with ADHD and ASD^27,34-37^. On the other hand, females diagnosed with ASD or developmental delay appear to have an increased burden of disruptive CNVs and rare, deleterious single nucleotide mutations compared with affected males^38-43^, though others have not found this effect for ultra-rare protein truncating-variants^11^. It is unknown whether rare CNVs are enriched in females with a broader range of NPs in a population sample.

Another possibility is that referral and diagnostic biases result in fewer females being diagnosed with NPs^44,45^. It has been suggested that childhood psychiatric problems in females either manifest differently than in males or are interpreted differently by parents and teachers, for example as anxiety or depression, instead of ADHD^44^. If this is the case, one might expect that genetic risk factors relevant to NPs would be more strongly associated with such other psychiatric problems in females. This possibility has yet to be examined.

In this study, we first set out to test whether rare CNVs are associated with NPs and anxiety/depression in a population sample of children. We used narrow (i.e. clinical diagnosis) and broad (i.e. screening measure) definitions of NPs to distinguish between clinically-recognised problems and sub-threshold population phenotypic variation. Second, we tested whether there are any sex differences in CNV burden in the context of NPs or anxiety/depression in the population.

## Method

### Sample description

The Child and Adolescent Twin Study in Sweden (CATSS) is a population study of twin children born in Sweden since July 1992^46^. The study started in 2004 and since then has been the method of inviting twins into the Swedish Twin Registry^47^. The cohort is being recruited when twins turn 9 years old but initially, parents of 12-year old twins were also invited to take part. Parents gave informed consent for study participation on behalf of their children. The study was approved by the Regional Ethical Review Board in Stockholm and Karolinska Institutet Ethical Review Board.

### Phenotypic measures

Information on five specific NPs (ADHD, ASD, motor problems, learning difficulties and tic problems) was obtained from two sources: registry-based diagnoses and screening via parental report. Diagnoses for these NPs, as well as anxiety and depression, were obtained through linkage with the National Patient Register (NPR) in Sweden, using each individual’s personal identification number. This NPR linkage contains information on all inpatient psychiatric care from 1987–2014 and outpatient consultations with specialists from 2001–2014. It includes best-estimate specialist diagnoses according to ICD-10 (International Classification of Diseases version 10) codes^48^. Study individuals were also screened for NPs using the Autism-Tics, ADHD, and Other Comorbidities inventory (A-TAC)^49^ administered to parents over the telephone when children were aged 9 or 12 years old. This measure has been reported to have good to excellent sensitivity and specificity for predicting clinical diagnoses, using validated clinical cut-offs^50^. These validated cut-offs were used to dichotomise children for each of the five NPs. See Table 1 for a list of ICD codes and details about the A-TAC measures used.

**Table 1:**
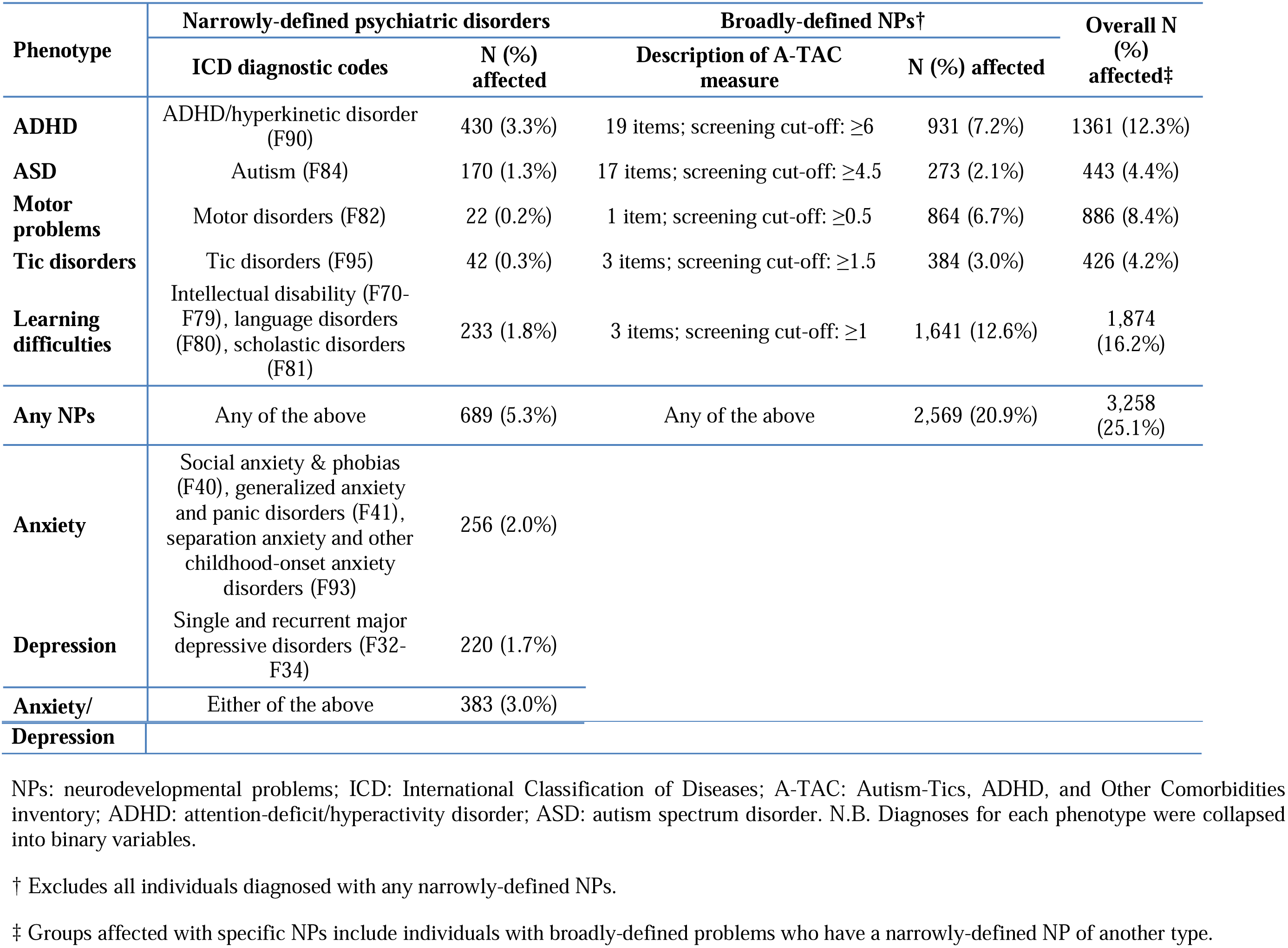
Phenotype descriptions

| Phenotype | Narrowly-defined psychiatric disorders |  | Broadly-defined NPs† |  | Overall N (%) affected‡ |
| --- | --- | --- | --- | --- | --- |
|  | ICD diagnostic codes | N (%) affected | Description of A-TAC measure | N (%) affected |  |
| **ADHD** | ADHD/hyperkinetic disorder (F90) | 430 (3.3%) | 19 items; screening cut-off: $\geq 6$ | 931 (7.2%) | 1361 (12.3%) |
| **ASD** | Autism (F84) | 170 (1.3%) | 17 items; screening cut-off: $\geq 4.5$ | 273 (2.1%) | 443 (4.4%) |
| **Motor problems** | Motor disorders (F82) | 22 (0.2%) | 1 item; screening cut-off: $\geq 0.5$ | 864 (6.7%) | 886 (8.4%) |
| **Tic disorders** | Tic disorders (F95) | 42 (0.3%) | 3 items; screening cut-off: $\geq 1.5$ | 384 (3.0%) | 426 (4.2%) |
| **Learning difficulties** | Intellectual disability (F70-F79), language disorders (F80), scholastic disorders (F81) | 233 (1.8%) | 3 items; screening cut-off: $\geq 1$ | 1,641 (12.6%) | 1,874 (16.2%) |
| **Any NPs** | Any of the above | 689 (5.3%) | Any of the above | 2,569 (20.9%) | 3,258 (25.1%) |
| **Anxiety** | Social anxiety & phobias (F40), generalized anxiety and panic disorders (F41), separation anxiety and other childhood-onset anxiety disorders (F93) | 256 (2.0%) |  |  |  |
| **Depression** | Single and recurrent major depressive disorders (F32-F34) | 220 (1.7%) |  |  |  |
| **Anxiety/** | Either of the above | 383 (3.0%) |  |  |  |

|  |
| --- |
| **Depression** |

NPs: neurodevelopmental problems; ICD: International Classification of Diseases; A-TAC: Autism-Tics, ADHD, and Other Comorbidities inventory; ADHD: attention-deficit/hyperactivity disorder; ASD: autism spectrum disorder. N.B. Diagnoses for each phenotype were collapsed into binary variables.
† Excludes all individuals diagnosed with any narrowly-defined NPs.
‡ Groups affected with specific NPs include individuals with broadly-defined problems who have a narrowly-defined NP of another type.

Because of substantial overlap across these 5 related phenotypes^50^, we derived two broad measures based on meeting criteria for one or more of these NPs. Individuals receiving any ICD diagnosis of NPs were classed as having “narrowly-defined NPs”, whereas individuals meeting any screening cut-off based on the A-TAC (but not receiving a clinical diagnosis) were classed as having “broadly-defined NPs”. Presence of NPs using these two definitions were the primary outcomes in analyses. Separate variables were also derived for each of the 5 specific NPs; see Table 1. Individuals who did not meet criteria for both broadly- and narrowly-defined NPs were considered as the unaffected/control group, for all analyses. A continuous measure of total NPs was derived by summing together the A-TAC scales for all 5 NPs (range of scores: 0–43). Specific NPs were categorized using a combined definition of either broadly- or narrowly-defined NPs, due to limited power (see Table 1).

### CNV calling and processing

DNA samples (from saliva) have been collected since 2008 from the participants after they are initially recruited to the study. Samples were genotyped using the Illumina Infinium PsychArray-24 BeadChip. For details of standard genotype quality control (QC) see Brikell et al^51^ and Supplemental Text. For details of CNV calling and QC, see Supplemental Text and for an overview see Figure S1. CNVs were called using the standard PennCNV protocol. Strict CNV- and sample-level QC was performed, following the protocol developed by the Psychiatric Genomics Consortium^20^. After all QC, the sample size for analysis consisted of N=12,982 individuals, including N=2,445 monozygotic and N=3,554 dizygotic twin pairs.

### CNV category definitions

CNV categories were defined based on size (all: >100kb; medium: 100-500kb; large: >500kb), type (deletion or duplication) and presumed relevance to NPs (relevant CNV, other CNV or no CNV). CNVs relevant to NPs were defined as those that either overlapped at least 50% with any of 69 genomic loci that have previously been implicated in ASD, intellectual disability and/or schizophrenia^8,20,52^ or with any gene within a set of 3,219 highly evolutionarily-constrained genes that are intolerant to mutations^53^. Further details about the definitions of these genomic regions can be found in the Supplementary Text. CNVs that overlapped with these loci are referred to as neuropsychiatric disorder/evolutionarily-constrained CNVs or ‘ND/EC-CNVs’ for short. For all categories of CNVs, the number of CNV segments was dichotomised to be ‘absent’ if no CNV call had been made for an individual within the category or ‘present’ if at least one called CNV met the criteria for that category.

### Data analyses

For the primary analysis, we examined the overall association of all CNVs passing QC, with narrowly- and broadly-defined NPs, as well as clinically-defined anxiety/depression. Several secondary tests were then performed to further examine the relationship between CNVs and psychiatric outcomes. First, CNVs were categorised by size and type to test the association of different CNV categories with each outcome. Next, we examined whether CNVs were associated with a continuous measure of NPs and whether significant associations were driven by specific NPs. Finally, we compared carriers of at least one ND/EC-CNV and carriers of other CNVs to controls and to each other, to further characterise the origin of significant associations.

Several analyses were performed to investigate sex-specific CNV effects. First, we tested for an overall difference in CNV burden by sex in the whole sample. To test the hypothesis that females with NPs are carriers of a higher burden of risk variants as compared to affected males, we compared CNV burden by sex in individuals with NPs. To test the alternative hypothesis that genetic risk factors relevant to NPs are more likely to be associated with anxiety/depression in females than in males, we compared CNV burden by sex in individuals with any anxiety or depression diagnosis. We also tested the impact of comorbid clinically-diagnosed NPs on this relationship by excluding individuals with narrowly-defined NPs and adjusting for the continuous measure of NPs. For all sex-specific analyses, males were coded as ‘0’ and females as ‘1’.

All analyses were performed using logistic or linear generalised estimating equations (with the package *drgee*^54^ in R-3.4.1), using family ID to cluster the data to account for related samples. The following covariates were included for all analyses: LRR SD, BAF SD, waviness factor, batch, and 5 principal components.

## Results

Table 1 shows details of the psychiatric outcomes. There was a strong association between a child receiving an ICD diagnosis of any NP (narrowly-defined NPs) and meeting study screening criteria for any NP (OR(CI)=9.66(8.04-11.59), p=1.2E-130). N=689 (5.3%) children had narrowly-defined NPs, N=2,569 children (19.8%) had broadly-defined NPs (based on meeting screening cut-offs, but not receiving clinical diagnoses), and N=9,724 (74.9%) children were unaffected by any definition of NPs. N=383 (3.0%) children had clinically-diagnosed anxiety and/or depression.

### Association of CNVs with psychiatric outcomes

There was an overall association between a child having any rare (<1% frequency) CNV >100kb in size and having any broadly-defined NPs (OR(CI)=1.13(1.03-1.25), p=0.014). There were no overall associations of presence of any CNVs with narrowly-defined NPs (OR(CI)=1.09(0.92-1.30), p=0.31) or anxiety/depression diagnoses (OR(CI)=1.15(0.92-1.42), p=0.22).

#### Secondary analyses

Associations between CNV categories of different sizes and types with NPs are shown in Figure 1a (see Table S1 for exact estimates). These secondary analyses indicated that medium-sized (100-500kb) deletions (OR(CI)=1.18(1.05-1.33), p=0.0053) and large (>500kb) duplications (OR(CI)=1.45(1.19-1.75), p=0.00017) showed the strongest associations with broadly-defined NPs. Although there was no overall association between CNVs and narrowly-defined NPs, large deletions showed a weak association (OR(CI)=1.85(1.14-3.01), p=0.013). There were no differences in presence of CNVs between children with narrowly- or broadly-defined NPs (p>0.05), except for large deletions (OR(CI)=1.82(1.06-3.11), p=0.028), which were more common in those with narrowly-defined NPs (Table S2). No CNV category was associated with anxiety/depression (p>0.05); see Table S3.

**Figure 1.**
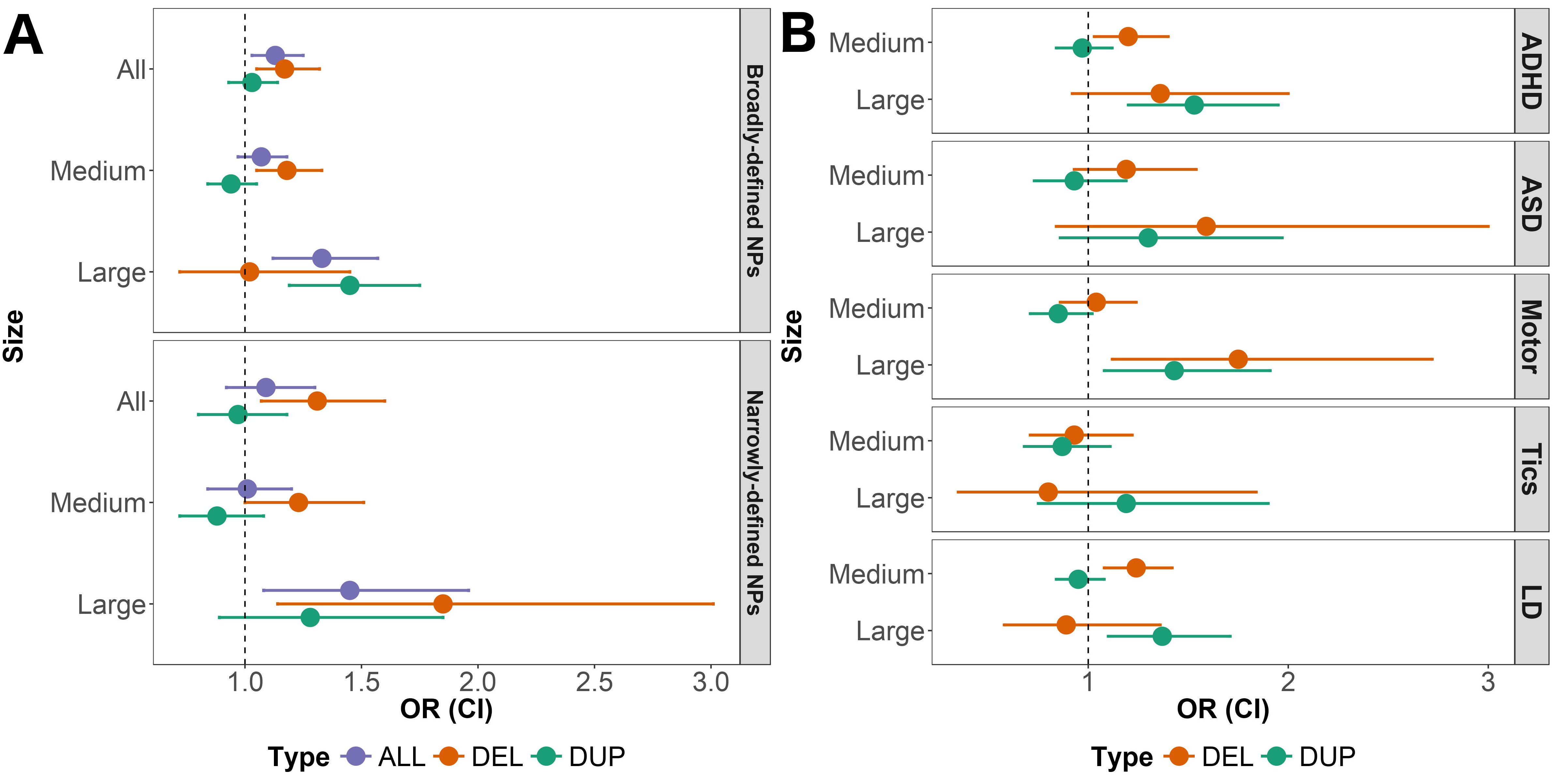
Association between presence of rare copy number variants (CNVs) categorized by size (all: >100kb; medium: 100-500kb; large: >500kb) and type (del: deletion; dup: duplication) with neurodevelopmental problems (NPs), defined as: a) broadly-defined NPs (i.e. meeting criteria using study screening measures but not receiving clinical diagnoses) or narrowly-defined NPs (i.e. receiving clinical diagnoses in the Swedish register data); b) specific NPs, using a combined definition of children meeting criteria for either broadly- or narrowly-defined NPs. A number of children are affected with more than one NP.
ADHD: attention deficit hyperactivity disorder, ASD: autism spectrum disorder; LD: learning difficulties; OR: odds ratio; CI: 95% confidence interval.

CNVs were also associated with the continuous measure of total NPs (Table S4), with the strongest associations once again observed for medium deletions (beta(SE)=0.30(0.11), p=0.009) and large duplications (beta(SE)=0.42(0.21), p=0.046), suggesting that a medium-sized CNV was associated with an average increase of 0.30 NP symptoms and a large duplication was associated with an average increase of 0.42 NP symptoms. Excluding individuals with clinically-diagnosed NPs reduced these effects (medium deletions: beta(SE)=0.26(0.10), p=0.0055; large duplications: beta(SE)=0.27(0.17), p=0.099).

Next, we examined the association of CNVs with specific NPs (see Figure 1b and Table S5). Medium deletions were significantly associated with ADHD (OR(CI)=1.20(1.03-1.40), p=0.017) and learning difficulties (OR(CI)=1.24(1.08-1.42), p=0.0017). Large duplications were associated with ADHD (OR(CI)=1.53(1.20-1.95), p=0.00051), motor problems (OR(CI)=1.43(1.08-1.91), p=0.013) and learning difficulties (OR(CI)=1.37(1.10-1.71), p=0.0047). Large deletions were also associated with motor problems (OR(CI)=1.75(1.12-2.72), p=0.013). Although the effect sizes for the association of CNVs with ASD and large duplications with tics were similar to the analyses of ADHD and learning difficulties, these results did not reach significance (p>0.05).

When analyses were limited to individuals who are carriers of ND/EC CNVs (i.e. CNVs likely to be relevant to NPs) as compared with control individuals, the observed effect sizes for large CNVs increased for both broadly-(OR(CI)=1.60(1.06-2.42), p=0.025) and narrowly-defined (OR(CI)=3.64(2.16-6.13), p=1.15E-6) NPs (Table S6). Figure 2 shows the proportion of NPs in the different CNV carrier groups. Carriers of other (non-ND/EC) large CNVs also had a higher proportion of broadly-defined NPs as compared with controls (OR(CI)=1.28(1.06-1.54), p=0.010). This association was non-significant for medium-sized CNVs and for narrowly-defined NPs as an outcome (p>0.05). On the other hand, carriers of large ND/EC CNVs had an increased risk of narrowly-defined NPs compared to carriers of other large CNVs (OR(CI)=3.65(1.83-7.27), p=2.3E-4); this effect was non-significant when examining medium-sized CNVs or broadly-defined NPs (p>0.05).

**Figure 2.**
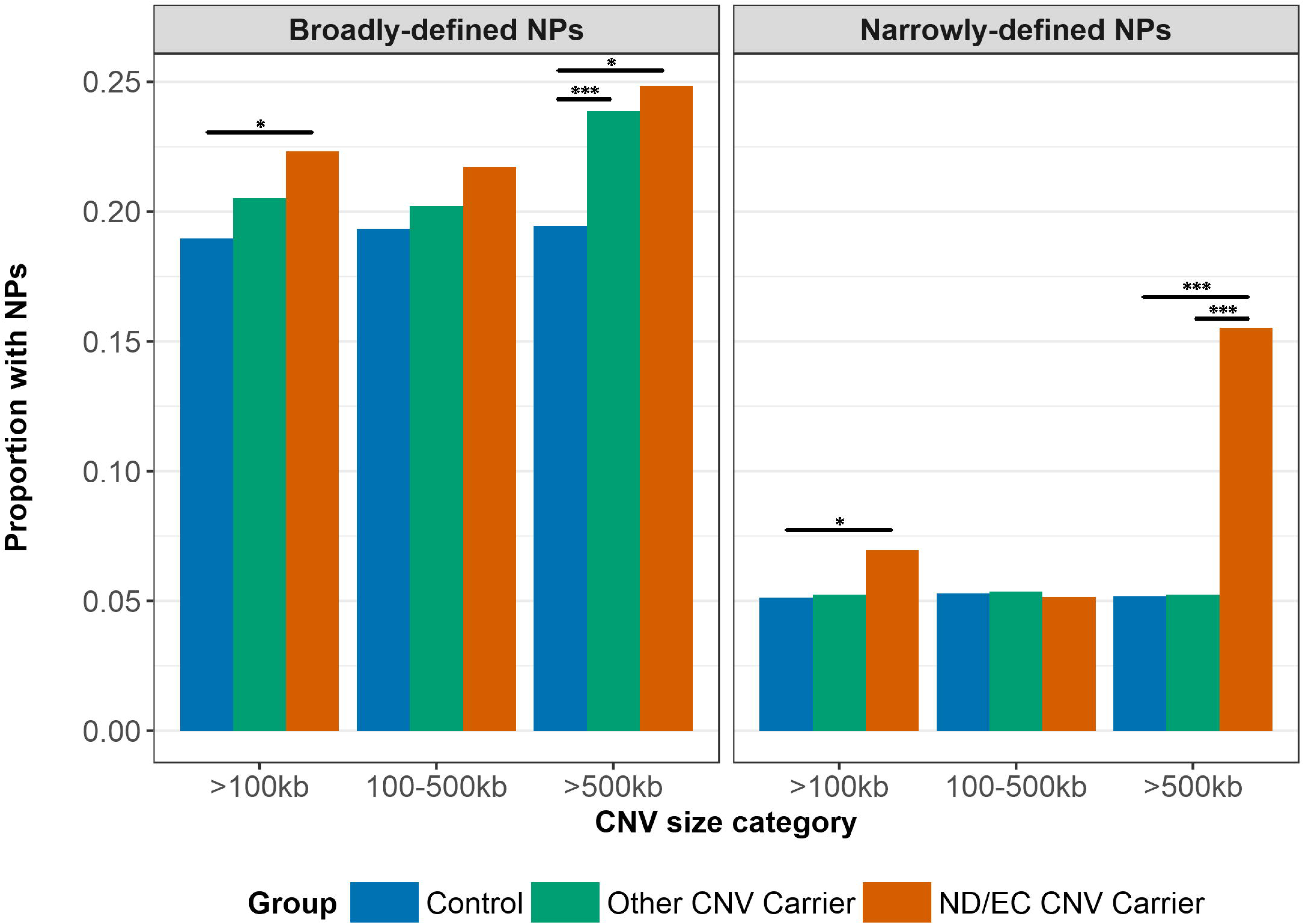
The proportion of broadly- and narrowly-defined neurodevelopmental problems (NPs) compared across carriers of copy number variants (CNVs) that affect neuropsychiatric disorders or evolutionarily-constrained genes (ND/EC CNVs), carriers of other CNVs and controls, with CNVs stratified by size. * p<0.05; ** p<0.01; *** p<0.001.

### Sex-specific analyses

N=1,486 (23.0%) males and N=1,083 (16.6%) females met criteria for broadly-defined NPs, N=482 (7.5%) males and N=207 (3.2%) females met criteria for narrowly-defined NPs and N=157 (2.4%) males and N=226 (3.5%) females met criteria for anxiety/depression; these sex differences were statistically significant [broad NPs: OR=0.63(0.57-0.69), p=7.4E-22; narrow NPs: OR=0.37(0.31-0.44), p=6.6E-27; anxiety/depression: OR=1.44(1.16-1.79), p=9.2E-4].

There was no overall difference in presence of CNVs by sex; see Table 2. There were also no significant sex differences in CNV presence in children with narrowly- or broadly-defined NPs. On the other hand, in those who had anxiety or depression diagnoses, females were significantly more likely than males to have large CNVs. The effect size for this association increased from 3.75 to 5.81 when individuals with comorbid narrowly-defined NPs were excluded. Adjusting for sub-threshold NPs by including the continuous measure of NPs as a covariate in this analysis further increased the effect size (OR=7.33(2.15-24.99), p=0.0015).

**Table 2:** Associations of CNVs with sex

| Sample | CNV Size | CNV presence (%) in females/males | OR(CI) | p |
| --- | --- | --- | --- | --- |
| **Whole sample (N=12,982)** | **All** | 43.9/44.4 | 0.99(0.91-1.08) | 0.85 |
|  | **Medium** | 39.7/40.3 | 0.99(0.91-1.07) | 0.77 |
|  | **Large** | 7.5/7.4 | 1.03(0.88-1.21) | 0.68 |
| **Broadly-defined NPs (N=2,569)** | **All** | 46.2/46.6 | 0.97(0.81-1.16) | 0.75 |
|  | **Medium** | 40.7/41.8 | 0.96(0.80-1.15) | 0.65 |
|  | **Large** | 9.4/8.7 | 1.06(0.78-1.44) | 0.73 |
| **Narrowly-defined NPs (N=689)** | **All** | 50.7/44.0 | 1.36(0.95-1.94) | 0.092 |
|  | **Medium** | 43.5/38.8 | 1.24(0.86-1.78) | 0.26 |
|  | **Large** | 10.1/9.5 | 1.18(0.62-2.22) | 0.61 |
| **Anxiety/Depression (N=383)** | **All** | 49.6/43.3 | 1.21(0.77-1.91) | 0.41 |
|  | **Medium** | 41.2/40.1 | 0.96(0.61-1.52) | 0.87 |
|  | **Large** | 12.4/3.8 | 3.75(1.45-9.68) | 0.0064 |
| **Anxiety/Depression without narrowly-defined NPs (N=269)** | **All** | 50.3/45.7 | 1.13(0.65-1.97) | 0.67 |
|  | **Medium** | 41.2/43.5 | 0.79(0.45-1.39) | 0.41 |
|  | **Large** | 13.6/3.3 | 5.81(1.70-19.89) | 0.0051 |

CI: 95% confidence interval; CNV: copy number variant; NPs: neurodevelopmental problems.

## Discussion

The results of this study add to the growing body of literature on the role of CNVs in psychiatric phenotypes, by demonstrating that rare CNVs are significantly associated with both narrowly- and more broadly-defined NPs in a general population sample of children. Our results are consistent with previous clinical studies of NPs, which reported significant enrichment of CNVs in individuals with clinically-diagnosed NPs^5-9^. In this study, we extend this knowledge by implicating CNVs in children with more broadly-defined, sub-threshold NPs. We found no association between rare CNVs and clinically-diagnosed anxiety and depression in this childhood sample, in contrast to a population study which found an association between neuropsychiatric CNVs and depression (though not anxiety) in adults^18^. Although we found no differences in CNV burden by sex in individuals with NPs, our results suggested that CNVs are enriched in females diagnosed with depression or anxiety, as compared to diagnosed males.

Several secondary analyses were used to characterise the role of CNVs in NPs and to pinpoint the categories of CNVs most relevant to NPs. Our results showed that narrowly-defined NPs (i.e. clinical diagnoses independent of the study design) have specific associations with particularly large (>500kb) CNVs, especially deletions and CNVs in regions previously implicated in other neuropsychiatric disorders and critical biological processes. Moreover, large deletions were enriched in narrowly-defined NPs compared to broadly-defined NPs. On the other hand, medium-sized (100-500kb) deletions and large duplications were associated with broadly-defined NPs. In addition to large CNVs that were likely to be relevant to NPs, large CNVs not affecting these regions were also relevant to broadly-defined NPs. Thus, the evidence suggests that especially highly deleterious CNVs are associated with clinical diagnoses of NPs, whereas arguably somewhat less deleterious CNVs are associated with NPs defined more broadly in the population.

This study suggests that rare genetic variants related to clinically-diagnosed disorders are also relevant to more broadly-defined problems in the general population, consistent with previous twin and common variant studies^12^. When examining NPs as a continuous distribution of symptoms, we saw a similar pattern of associations to when NPs were defined dichotomously, with weaker effects when clinically-diagnosed individuals were excluded. These results support the idea that neurodevelopmental disorders reflect the quantitative extreme of genetic factors operating dimensionally among individuals in the general population.

Information on multiple different NPs was deliberately combined in this study because of the high comorbidity and shared genetic risk between different disorders^1,55^. It has been suggested that a broad and heritable general psychopathology factor underlies NPs^56,57^. Our current results lend support to this idea by demonstrating that CNVs are to a large degree nonspecific, affecting multiple related neurodevelopmental outcomes in children from the general population. Although the analyses of specific NPs suggested that these classes of CNVs were most strongly associated with ADHD, motor problems and learning difficulties, these outcomes were in fact the most common ones in the sample (see Table S1); the estimated confidence intervals overlapped to a large degree for the different NPs and it is likely that these comparisons were affected by decreased power for ASD and tic problems.

The second aim of our study was to determine whether CNVs showed sex-specific associations that could elucidate the excessive male bias seen in NPs. In contrast to previous studies which have found an increased burden of large, rare CNVs in females with ASD and developmental delay^38,39,42^, our results showed no differences in CNV presence in males and females with NPs. This inconsistency may be due to ascertainment and measurement differences. The previous studies were clinically-ascertained samples of children meeting research-based diagnostic criteria for NPs, whereas the current dataset is a childhood population cohort study, with NPs defined using either a lifetime clinician’s diagnosis (that is independent of study recruitment) or a broad study screening measure.

Interestingly, we detected more large CNVs in females with ICD diagnoses of anxiety or depression than in diagnosed males. This result could suggest that large, rare CNVs manifest differently in males and females. Adjusting for comorbid NPs (by excluding individuals with narrowly-defined NPs and including sub-threshold NP traits as a covariate) strengthened the association. This indicates that comorbid NPs in females with anxiety or depression do not explain the observed results. One interpretation is that large, rare CNVs may be more likely to be associated with anxiety or depression diagnoses in females. A complementary interpretation is that males with such CNVs may be less likely to be diagnosed with anxiety or depression; although the rate of large CNVs was similar by sex in the whole sample (females: 7.5%; males: 7.4%), in individuals with diagnoses of anxiety or depression but not NPs, females had an increased rate (13.6%) whereas males had a decreased rate (3.3%) of large CNVs. In contrast to the male bias seen in NPs, epidemiological studies show a female bias towards diagnoses of anxiety and depression^58,59^. Our results hint that this prevalence difference may be in part due to sex-specific referral or diagnostic biases, where female CNV carriers who manifest psychopathology and are referred for clinical assessment may be more likely to receive diagnoses of anxiety or depression rather than NPs, which are more common in males. However, the sample size of diagnosed individuals is limited in the current data and larger studies are needed to replicate and further interpret this preliminary finding.

The strengths of this study include the use of a large homogenous population sample with genome-wide data as well as multiple assessments of NPs (based on study screening using parental questionnaires and study-independent ‘real-life’ clinician’s diagnoses). However, as this was a general population cohort, the sample size of individuals with clinical diagnoses was limited, thus affecting the power of the analyses. The power was further limited for stratifying rare CNVs by size, type and relevance to NPs so we are unable to draw conclusions about the specific loci (and for the sex-specific analyses even the category of CNVs) that may be driving observed associations. Another limitation is that certain types of CNVs and other structural variants (e.g. individuals with possible aneuploidies) were excluded from the study during quality control, which may have affected the generalisability of our results to a broader class of rare structural variation in the general population.

Future studies using large genome-wide datasets are needed to further determine the role of specific rare genomic loci in NPs and other psychiatric phenotypes in the general population. Our results add to studies of CNVs in adult populations^16-18^, demonstrating that population datasets with information on broad psychiatric problems will be an important addition to case-control studies in characterising the role of rare variation in relation to psychiatric phenotypes.

Our study also finds evidence of a novel association between large, rare CNVs with anxiety and depression in females compared to males. This points to a possible role of sex-specific manifestation of CNVs that would benefit from further study.

## Acknowledgements

Dr Martin was supported by the Wellcome Trust (Grant No: 106047). Karolinska Institutet and the Swedish Research Council financially support the Swedish Twin Registry, which the CATSS dataset is part of. CATSS also has support from the Swedish Research Council for Health, Working Life and Welfare. Funders were not directly involved in the study.

